# The fitness of an introgressing haplotype

**DOI:** 10.1101/2022.09.27.507129

**Authors:** Andrius J. Dagilis, Daniel R. Matute

## Abstract

The genomic era has made clear that introgression, or the movement of genetic material between species, is a common feature of evolution. Examples of both adaptive and deleterious introgression exist in a variety of systems. What is unclear is how the fitness of an introgressing haplotype changes as species diverge, or as the size of the introgressing haplotype changes. In a simple model, we show that early in the process of divergence, introgression of large haplotypes can be favored more than introgression of individual alleles. The key insight is that alleles from a shared genetic background are likely to have positive epistatic interactions, increasing the fitness of a larger introgressing block. The buildup of incompatibilities between diverging species in the form of deleterious epistasis eventually favors the introgression of small haplotypes as the number of diverged alleles increases, and eventually even single alleles with positive direct effects can be selected against. This model is consistent with observations of a positive relationship between recombination rate and introgression frequency across the genome, however it generates several novel predictions. First, the model suggests that the relationship between recombination rate and introgression may not exist, or may be negative, in recently diverged species pairs. Furthermore, the model suggests that introgression that replaces existing derived variation will always be more deleterious than introgression at sites carrying ancestral variants. These predictions are tested in an example of introgression in *D. melanogaster*, with some support for both.

## Introduction

Before reproductive isolation is complete between two taxa, these taxa are often able to exchange migrants, form hybrids, and as a result exchange genes (for a recent review, see Edelman and Mallet (2021)). Even a small number of hybrids between young species pairs can lead to introgression of genetic material between them (Barton and Bengtsson 1986; Barton 2001), and studies across Eukaryotes have identified introgression between a vast range of species (Dagilis et al. 2022). Studies of hybrids have identified that the degree of divergence between the parental species strongly predicts hybrid fitness (Coyne and Orr 2004; Matute et al. 2010; Moyle and Nakazato 2010; Coughlan and Matute 2020), and many models have considered how the fitness of these hybrids changes over the course of divergence, whether using insights from the build-up of incompatibilities (Orr 1995; Turelli and Orr 2000; Gavrilets 2004; Dagilis et al. 2019) or through Fisher’s Geometric Model (Barton 2001; Fraïsse et al. 2016; Simon et al. 2018). Generally, these models find that there is a “grey zone” of speciation - when hybrid fitness is reduced, yet remains high enough that hybrids between the two species survive and mate sufficiently to enable gene flow between the species (Roux et al. 2016). There are many fewer models of introgression, and those that exist generally assume that introgression is primarily deleterious (Harris and Nielsen 2016; Martin and Jiggins 2017; Petr et al. 2019; Veller et al. 2021; Pfennig and Lachance 2022), however we know that sometimes introgression is adaptive (Huerta-Sánchez et al. 2014; Norris et al. 2015; Edelman and Mallet 2021), and so what types of loci and what fraction of the genome in general can introgress remains unknown, but it is likely that introgressed regions are rarely neutral.

Several patterns are suggestive of the non-neutrality of introgressed regions. Empirical studies have observed a relationship between local recombination rate and introgression (Muirhead and Presgraves 2016; Schumer et al. 2017; Ravinet et al. 2018; Silva et al. 2018; Edelman et al. 2019). This relationship has been explained by “hybrid load”, or the idea that alleles from a minor parent ancestry (the ancestry with less representation in the hybrid) are selected against (Schumer et al. 2016; Veller et al. 2021). The mechanism for this selection is less clear - it is possible that the introgressing haplotypes carry weak deleterious epistatic interactions with the receiving genome (e.g., swordtail fish (Schumer et al. 2017), or butterflies (Martin et al. 2019)), or that they are simply directly deleterious due to having evolved in a more inbred population (e.g., Neanderthals (Harris and Nielsen 2016; Juric et al. 2016; Petr et al. 2019)). In both cases a clear negative relationship between the size of the introgressing haplotype and its fitness is expected. By contrast, a negative relationship between introgressed ancestry and recombination has been observed in *D. melanogaster* (Pool 2015). Recent work has used simulations to demonstrate that this relationship could be in part explained by positive selection on introgressed regions (Duranton and Pool 2022). Empirical evidence of positive selection of introgressed alleles is extensive (see Edelman and Mallet (2021) for a review).

Positive effects of introgression have demonstrated how heterotic effects resulting from overdominance can increase the probability of neutral markers introgressing (Pfennig and Lachance 2022) even when deleterious epistasis with the host genome is frequent. However, introgressing haplotypes are likely to carry epistatic interactions not only with the receiving genome, but also among the introgressing alleles as well. Thus, larger introgressing haplotypes may in fact be buffered against weak negative epistatic effects by carrying co-adapted sets of alleles. These effects are particularly relevant when recombination cannot break up the haplotype for some reason – whether it is due to the lack of recombination in general, such as horizontal gene transfer in asexual lineages, or due to suppressed recombination along sex determining regions (Dixon et al. 2019), inversions (Fontaine et al. 2015; Norris et al. 2015) or other methods (Poelstra et al. 2014). Currently, no theoretical approach has examined how the fitness of an introgressing haplotype changes depending on both its size and divergence between the two populations.

In this paper, we fill that theoretical gap. We begin by asking what selective forces act on different types of introgressing alleles - novel alleles which introduce derived (from the ancestor) variants at an ancestral site, ancestral alleles which introduce an ancestral variant at a derived site in the host population, and replacement alleles which introduce derived variants at derived sites. We then ask how the fitness of small haplotypes consisting of a small number of introgressing alleles changes as the total number of substitutions in the host population increases. We find that the overall divergence between species is a key predictor of the types of haplotypes that can successfully introgress. In general, introgressing ancestral variants are more strongly selected against than novel derived alleles, suggesting that regions that have diverged rapidly may be more resilient to introgression. Early in divergence, large haplotypes carrying many derived alleles may make it across species barriers with relative ease. As divergence continues, larger haplotypes are more and more strongly selected against, but primarily due to their effects in breaking up existing epistatic interactions in the receiving population, and not due to novel incompatibilities that they might carry. This model creates a new framework with which introgression over the course of speciation can be explained, provides several testable predictions, and suggests further avenues for both theoretical and empirical approaches. We examine some of these predictions in genomic data from populations of *D. melanogaster* which have experienced recent introgression (Pool 2015; Coughlan et al. 2021) and find that introgression is primarily detected in regions which show little divergence from a common ancestor in the host population but are highly diverged in the source of introgression.

## The Model

We model a haplotype introgressing from a source population (A) into a host population (B). We assume a total of *b* derived substitutions have been fixed in B since it diverged from A (Figure 1A). We ask what the fitness of a rare introgressing haplotype is (*W*_I_) compared to the average individual in the host population 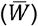, and calculate the effective selection coefficient (*σ*_*I*_) for the haplotype, equal to 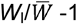. We will consider both direct and digenic epistatic fitness effects among alleles. We assume that all fitness effects are multiplicative, but our approximations are congruent to additive fitness. We will first examine results in a haploid population, but they extend readily to diploid cases. First, let’s examine the fitness of the host population (compared to the ancestor). This fitness will depend on 𝔹, the set of substitutions fixed in that population, for a total of *b* substitutions, and the average direct and epistatic fitness effects:

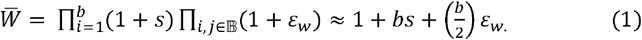

Where *s* is the average direct fitness effect of substitutions, and 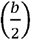 denotes the binomial coefficient “*b* choose 2” equal to the number of pairs among *b* total substitutions. Finally, *ε*_*W*_ denotes the strength of epistatic interactions of alleles that have evolved “within” the same genetic background. The approximation on the right side of equation (1) is accurate whenever the selective effects (both direct and epistatic) are very small and have low variance. The exact form of equation (1) is used to calculate all results in figures, but the approximation is useful in describing how parameters impact fitness. For instance, note that fitness in the host population depends linearly on direct fitness, but at least quadratically on epistatic effects.

**Figure 1:**
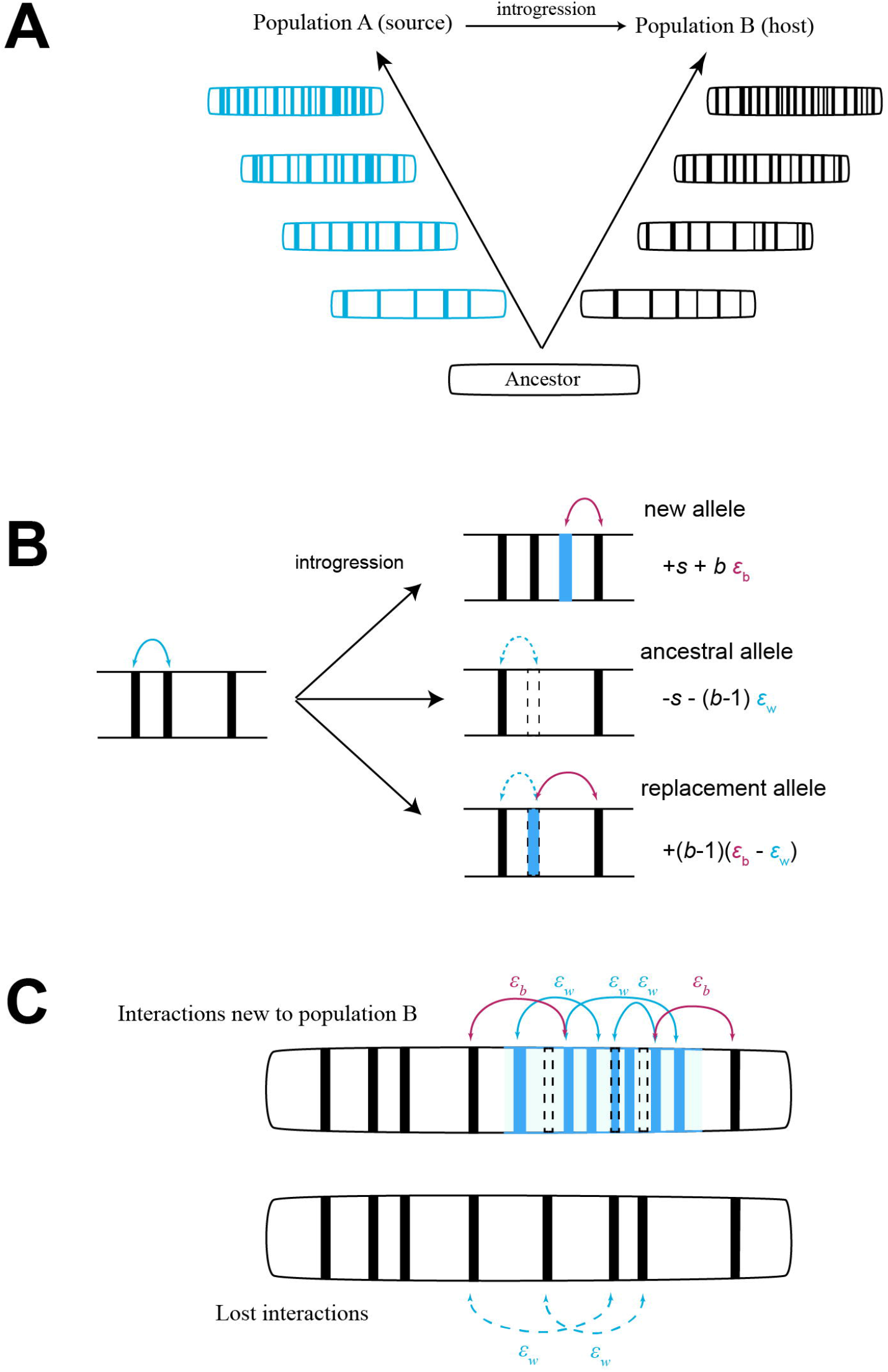
Outline of model assumptions. **(A)** Two populations (A and B) evolve independently, fixing substitutions in their genome, represented by filled bars, until introgression at some point moves alleles from A into B. **(B)** Three types of alleles may introgress as a result: new alleles which are derived in A and ancestral in B, ancestral alleles which are ancestral in A and derived in B and replacement alleles in which both A and B have fixed alternate variants. Each of these types of alleles will on average have different fitness effects. **(C)** An introgressing haplotype of length *x* carrying all three types of alleles will both introduce novel epistatic interactions (solid arrows), and remove existing ones (dashed) in individuals carrying the haplotype compared to others in the population.

### Selection on individual alleles

We can now consider introgression of three different classes of alleles (Figure 1 B). The first is a substitution which occurred in population A but not in B, termed “new” alleles. The selection coefficient on this allele can be expressed as:

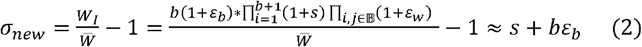

That is, the allele carries one new direct fitness effect, as well a *b* epistatic interactions of strength *ε*_*b*_ – epistasis of alleles “between” populations. Unlike within population epistatic interactions, these will be epistatic interactions that have not been tested by selection (as they are between alleles evolved in different backgrounds) and so are likely to be quite different (see Dagilis et al. (2019) for simulations confirming this intuition). Indeed, while *ε*_*W*_ can on average be expected to be positive, *ε*_*b*_ is likely to be both negative and an order of magnitude weaker. What these parameters suggest is that even alleles with directly beneficial fitness effects will be selected against as populations diverge (*b* increases).

A second type of allele that can introgress is what we will call an ancestral allele – a position in the genome in which the host population has fixed a substitution, but the introgressing allele carries an ancestral variant. Following the logic from above, the selection coefficient of an allele of this type can be expressed as:

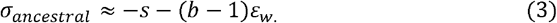

In this case, the introgression of the ancestral allele removes a direct fitness effect as well as *b*-1 epistatic effects with existing substitutions. Thus, it is immediately clear that fitness effects of introgressing ancestral variation will in general be more deleterious than those of introgressing new alleles under expected parameter space, since they remove whatever the direct fitness effect is and remove positive epistasis. As with the introgression of new alleles, these become increasingly deleterious throughout divergence.

Finally, we can consider what occurs when an introgressing allele is derived compared to the ancestral allele, but also replaces an existing derived allele in population B. In this case, it is reasonable to assume that the two fitness effects are essentially combined – we can call this type of allele a “replacement” allele and express its selection coefficient as:

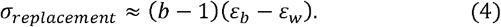

Here again, whenever *ε*_*b*_ < 0 *and ε*_*w*_ > 0, increased divergence leads to stronger and stronger selection against introgression, however direct selection coefficients do not play a role. Since epistasis may often be orders of magnitude weaker than direct fitness effects, these types of alleles may be very weakly selected while *b* is small (early in divergence). Note that in the special case *ε*_*b*_ = *ε*_*w*_, no selection against these types of alleles is expected.

When more than a single allele introgresses, the effect of the full haplotype is not simply a sum of the above effects, but also includes effects between the alleles on the haplotype itself. The only new type of interaction will be within population epistatic interactions (*ε*_*w*_) between alleles coming from population A (either new or replacement alleles). That is, each pair of those alleles has its own small epistatic interaction of strength *ε*_*w*_.

### Selection on haplotypes

Introgression of a larger block of alleles is likely to contain loci of all three types (Figure 1C). To simplify notation somewhat, we can consider that the introgressing haplotype will carry alleles at two types of sites – ones in which the source population (A) has fixed substitutions, and ones in which the host population (B) has fixed substitutions (with replacement alleles being alleles of both types). Letting the numbers of these be *x*_A_ and *x*_B_ respectively, we define the size of the haplotype (*x*) as the total number of alleles it is carrying, which will be equal to the number of new, ancestral and replacement sites. Since sites with derived alleles fixed in A can be either new or replacement alleles, *x*_A_ = *x*_new_ + *x*_replacement_ and *x*_B_ = *x*_ancestral_ + *x*_replacement_. With all of these assumptions, the effective selection coefficient of an introgressing haplotype in a haploid population can be expressed as:

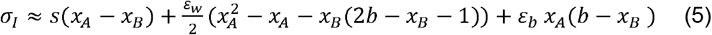

We can further simplify the above relationship by assuming that replacement alleles will be quite rare early in the process of divergence, and letting the fraction of *x* which is ancestral alleles be denoted as *f* with the remainder being new alleles (e.g. *fx = x*_ancestral_, (1-*f*)*x* = *x*_new_). The selection coefficient on an introgressing haplotype can then be expressed as:

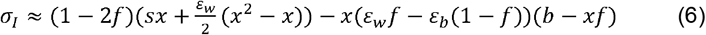

Note that divergence between populations (*b*) only appears as a linear term, while the size of the introgressing haplotype (*x*) appears as a quadratic term. As a result, the size of the haplotype can have a larger effect on its fitness than the number of substitutions fixed in the host population.

We can parameterize the exact form of Equation (5) to examine the exact effects of *b* and x. We assume that *ε*_*b*_ = -10^−4^ and that *ε*_*w*_ is an order of magnitude larger and positive. When *b* is small, introgression of large haplotypes carrying new alleles is strongly favored (Figure 2A). As divergence continues, the size of the introgressing haplotype necessary to overcome the existing incompatibilities becomes larger, but counter-intuitively, small haplotypes are still strongly selected against (Figure 2A). A second effect of divergence is that haplotypes become increasingly unlikely to carry solely new alleles. Introducing ancestral variants decreases the fitness of the introgressing haplotype strongly (Figure 2B). Under parameters in which a haplotype carrying 30 novel alleles provides a nearly 20% increase in fitness compared to the host population, if just a quarter of the alleles are instead ancestral, the haplotype is equally selected against. If we further expand to vary the number of new, ancestral and replacement alleles at the same time, we find that replacement alleles act largely like ancestral alleles, especially later in divergence (Figure 3). While we choose very large epistatic effects here for illustrative purposes, they are not outside the realm of observed epistatic effects in yeast mutants (Costanzo et al. 2016). Smaller epistatic effects are likely to change the scale of divergence needed to see strong selection against introgressing haplotypes as well as the size of the introgressing block before it becomes positively selected. We can predict both of these quantities by examining the inflection point of Equation (6).

**Figure 2.**
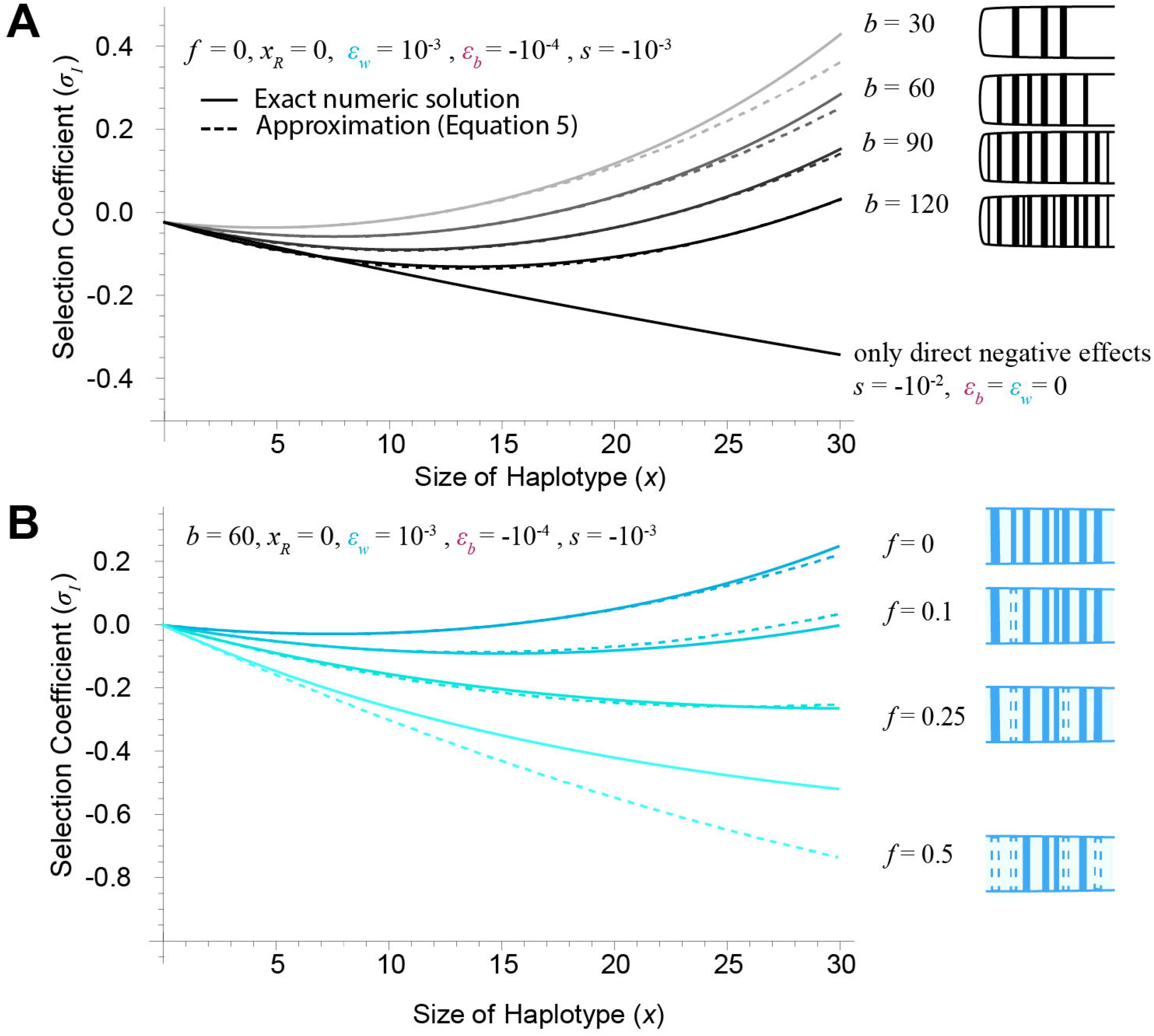
Key results. **(A)** When divergence is low, even fairly small introgressing haplotypes carry sufficient positive interactions between each other to cancel out potential deleterious effects. As divergence increases, however, the fitness of smaller haplotypes rapidly declines, and it takes larger and larger haplotypes to cancel out potential DMIs. If each allele has only direct negative selection on it in the receiving population (dashed line) a simple linear relationship in fitness is expected. Note that we assume direct selection on each allele is on average negative, and so the positive selection is entirely driven by epistasis. **(B)** When the introgressing haplotype is only carrying novel alleles, it is able to move in easily. However, as it replaces more and more of existing substitutions in the receiving population, it becomes more and more strongly selected against. In both panels, it is assumed all alleles are carrying either new or ancestral variants, with no replacement alleles. Solid lines show exact numeric solutions, while dashed lines show approximations from Equation (5).

**Figure 3:**
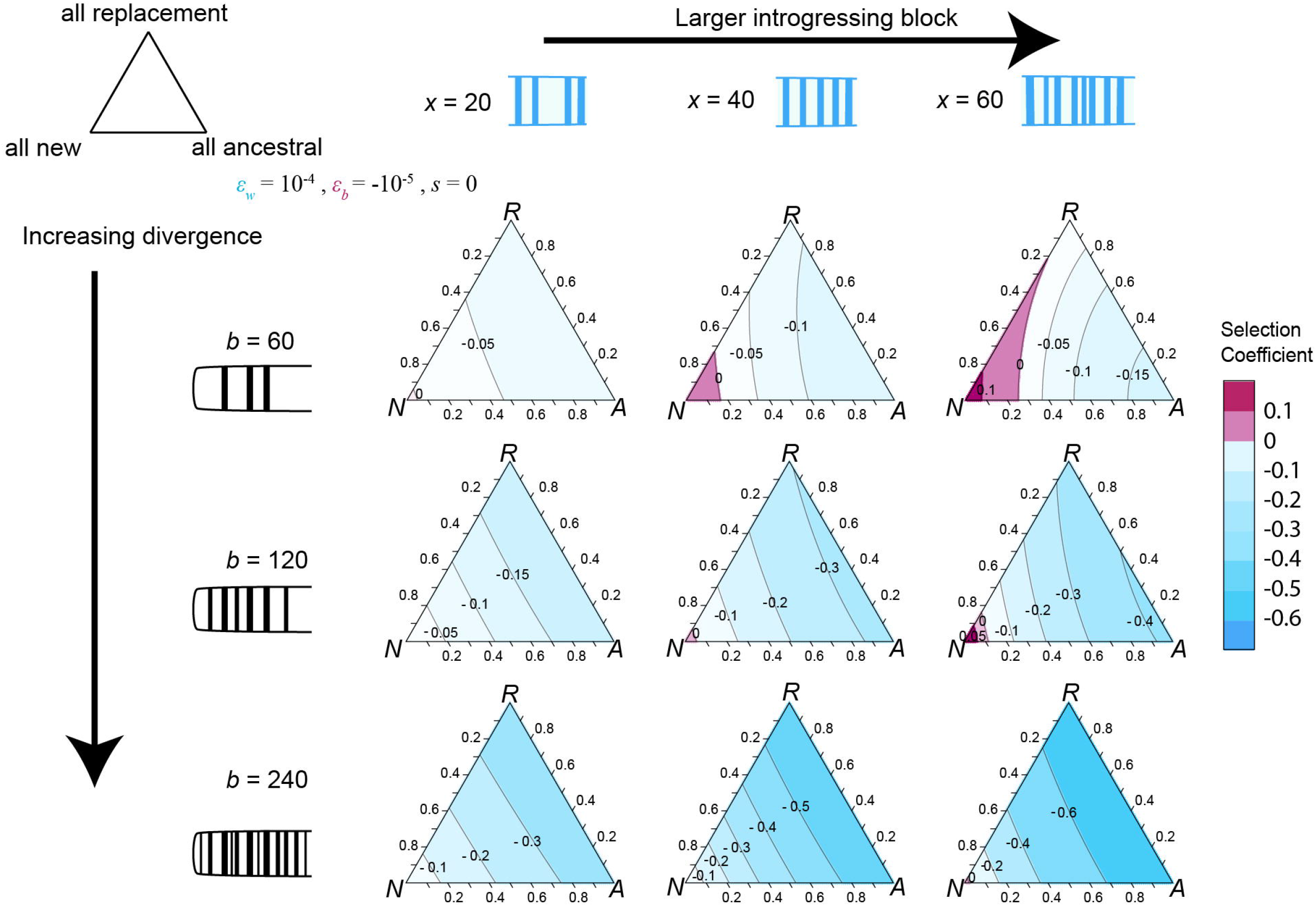
The effect of replacement alleles mimics introgressing ancestral variation. Examining how the selection coefficient on an introgressing haplotype changes as the fraction of substitutions it carries varies between new, ancestral and replacement alleles. Parameters as in Figure 2 () and calculations based on exact numerical evaluation of fitness (see Supplementary Figure X for approximation using Equation 5). Early in divergence (top row), carrying more ancestral variants has a more strongly deleterious effect than carrying replacement alleles. However, as divergence increases (lower plots), haplotypes that carry replacement alleles begin looking largely like haplotypes which carry entirely ancestral variants. In all cases, haplotypes carrying only novel alleles are most strongly beneficial.

Since *σ*_*I*_ = 0 when *x* = 0 (by definition), the quadratic shape of the fitness function (Equation (6)) and reasonable parameter assumptions (*ε*_*w*_ > 0 > *ε*_*b*_) suggest there is a minimum fitness of an introgressing haplotype dependent on its size (Supplementary Figure 1). This minimum will occur when the derivative of Equation (6) with respect to x is 0. If we ignore direct fitness effects (i.e. let *s* = 0), this value occurs at:

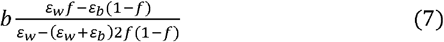

When *f* = 0, i.e. only new alleles are being introduced this is simply 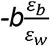, so it depends entirely on the ratio of between and within population epistasis (Supplementary Figure 1). If between population epistasis is several orders of magnitude weaker than within, the most strongly selected against haplotypes will be a small fraction of the total divergence between populations. Any larger haplotypes will be increasingly less deleterious, and potentially positively selected. When both *ε*_*b*_ and *ε*_*W*_ are positive, this inflection point is negative meaning all introgression is positively selected. As within and between epistasis become more similar, this value gets closer and closer to *b*, meaning larger haplotypes are necessary to overcome any deleterious effects. When *f*=1/2 (so half of the alleles are replacing existing variants) the above is always equal to *b*, meaning that haplotypes are increasingly selected against until they replace more than half of the existing substitutions, and introduce an equal amount of new alleles. Finally, when *f*=1, Equation (7) is again equal to *b*, meaning that haplotypes that are only replacing existing variants are always more strongly selected against as they become larger (Supplementary Figure 1).

### Selection in diploids

While results are straightforward for the haploid model, several new parameters need to be introduced for a diploid model. Direct fitness effects are assumed to have dominance *h*. Epistatic interactions may also have varying degrees of dominance (Supplementary Figures 2,3). In particular, epistatic interactions may occur at different strengths depending on whether both derived alleles are heterozygous, one is heterozygous and the other is homozygous, or both are homozygous (Turelli and Orr 2000; Dagilis et al. 2019). We parameterize this by setting the interaction strength when both alleles are homozygous as ε, while homozygous-heterozygous interactions are assumed to be on average of strength *a*_2_*ε* and heterozygous-heterozygous interactions are of strength *a*_1_*ε*. A closed form expression for approximate fitness of a haplotype introgressing in a diploid population can be found in materials and methods (Equation (8)), but the addition of two extra parameters makes simple analytical conclusions more difficult to draw. Assuming that *a*_1_ and *a*_2_ are roughly proportional, several broad patterns can be identified. As epistasis becomes recessive, the role of new alleles is strongly weakened (Supplementary Figure 3). Especially early in the process of introgression, the introgressing haplotype is almost always going to be found in heterozygotes. As a result, positive interactions between alleles it carries will not be fully expressed, while the loss of positive epistasis from any ancestral/replacement alleles will be very strong (Supplementary Figure 3). On the other hand, dominant epistasis would suggest that positive epistatic effects among introgressing alleles are expressed nearly as strongly as in the haploid case, without suffering loss of positive effects from ancestral/replacement alleles (Supplementary Figure 3). If epistasis tends to be dominant, therefore, introgression should on average be more positively selected than if it is recessive. In nature, of course, introgression is likely to vary in dominance broadly both within and between systems.

### Effects of recombination

So far, we have discussed the model in terms of haplotypes unaffected by recombination. This simplifying assumption allows us to infer the effective selection coefficient of a set of genes, but is not likely to be held in a variety of cases of introgression. We were unable to extend the results to an arbitrary linkage map, however, we can make some conclusions about introgression across the genome based on the results from the recombination free approach. Imagine the genome is split up into haplotypes, the size of which is inversely correlated to local recombination rates. In turn, we would expect regions of the genome with high recombination to have small introgressing haplotypes, while regions of the genome with low recombination may carry much larger introgressing blocks. To further simplify, we will assume that any individual is at most carrying a single region with introgressed alleles. While the average fitness in the population (equation 1) will change with many introgressing haplotypes of various sizes, their relative fitness should still show the same shape as average population fitness only appears in the denominator. As a result, we can make some predictions – early in the process of divergence, large introgressing blocks (found most likely in regions of low recombination) will have higher fitness than small ones (found in regions of high recombination). Thus, there may be a negative relationship between local recombination rate and introgression. Later in divergence, large haplotypes are less fit than small ones, and so a positive relationship between recombination rate and introgression is expected to occur. The latter pattern has been seen in many studies of introgression between species (Schumer et al. 2017; Ravinet et al. 2018; Martin et al. 2019), while a negative pattern between recombination and introgression was observed in gene flow between populations of *Drosophila melanogaster* (Pool 2015).

### Introgression in *Drosophila melanogaster*

Within Africa, *D. melanogaster* is strongly genetically structured, with multiple distinct ancestries geographically distributed throughout the region (Coughlan et al. 2021). Subsequent gene flow between these ancestries has been detected in multiple studies (Pool 2015; Coughlan et al. 2021). A particularly intriguing case is introgression of African ancestry into North American populations, in which a negative relationship between recombination rate and introgressed ancestry was previously identified (Pool 2015; Duranton and Pool 2022). In previous work, we identified recent introgression between an ancestry primarily present in Western Africa (called West) and an Out of Africa ancestry primarily present among North American flies (referred to as OOA2), confirming prior results. This introgression event represents an opportunity to test some of the predictions of our model. In previous work, we had calculated a measure of introgression proportion within local genomic windows (*f*_D_) as well as population differentiation (*F*_ST_) in non-overlapping 200 SNP windows across the genome for these populations (Coughlan et al. 2021). Here, we use that data to test several predictions of our model.

We first calculated the Population Branch Statistic (PBS) (Yi et al. 2010) for each of three populations: OOA2, West and South1 – the latter being an ancestry present in Southern Africa with no evidence of introgression with either of the other two populations. We use *F*_ST_ calculated excluding any individuals that carried more than 1% introgressed ancestry according to ancestry estimates performed using PCAngsd (Meisner and Albrechtsen 2018). We next excluded all windows in which PBS was negative for any of the populations as a negative value of PBS is not intuitive to interpret (however, all results hold even when these windows are retained, Supplementary Figure 4), and the resulting values naturally sum to 1, letting us ask what proportion of evolution has likely occurred on each branch among the three populations. A PBS_West_ value of 1 for a window would therefore indicate that all substitutions within that window have taken place along the branch to the Western population, as South1 serves as an outgroup and likely represent ancestral variants. We then plotted the values of *f*_D_ across the genome on a ternary plot for all three values of PBS (Figure 4). While the majority of our windows demonstrated large PBS values for the OOA2 population, in line with this population having diverged as *D. melanogaster* expanded into North America, windows with larger *f*_D_ values were more broadly distributed, with many windows clustering near the largest PBS values for the West population we could identify (Figure 4B). This recent introgression event seems to confirm some of the predictions made in our model, as these populations with low divergence show overall a negative relationship between introgression and recombination (Figure 4C) as observed previously (Pool 2015; Duranton and Pool 2022) and windows that have shown higher divergence in the host population show overall resilience to introgression (Figure 4D), with both relationships significant in a linear model of the data (*F* values of 98.8, 2583, 0.004 for recombination, PBS_OOA2_ and their interaction respectively in an ANOVA). Note that because the relative branch scores are calculated using individuals with no admixed ancestry, this pattern is not driven by introgression reducing PBS scores for the OOA2 population. Nonetheless, very few windows in our data set had large PBS scores for the West population, and as a result one of the key predictions of our model – that regions of the genome carrying entirely novel variation should introgress very easily, remains untested.

**Figure 4:**
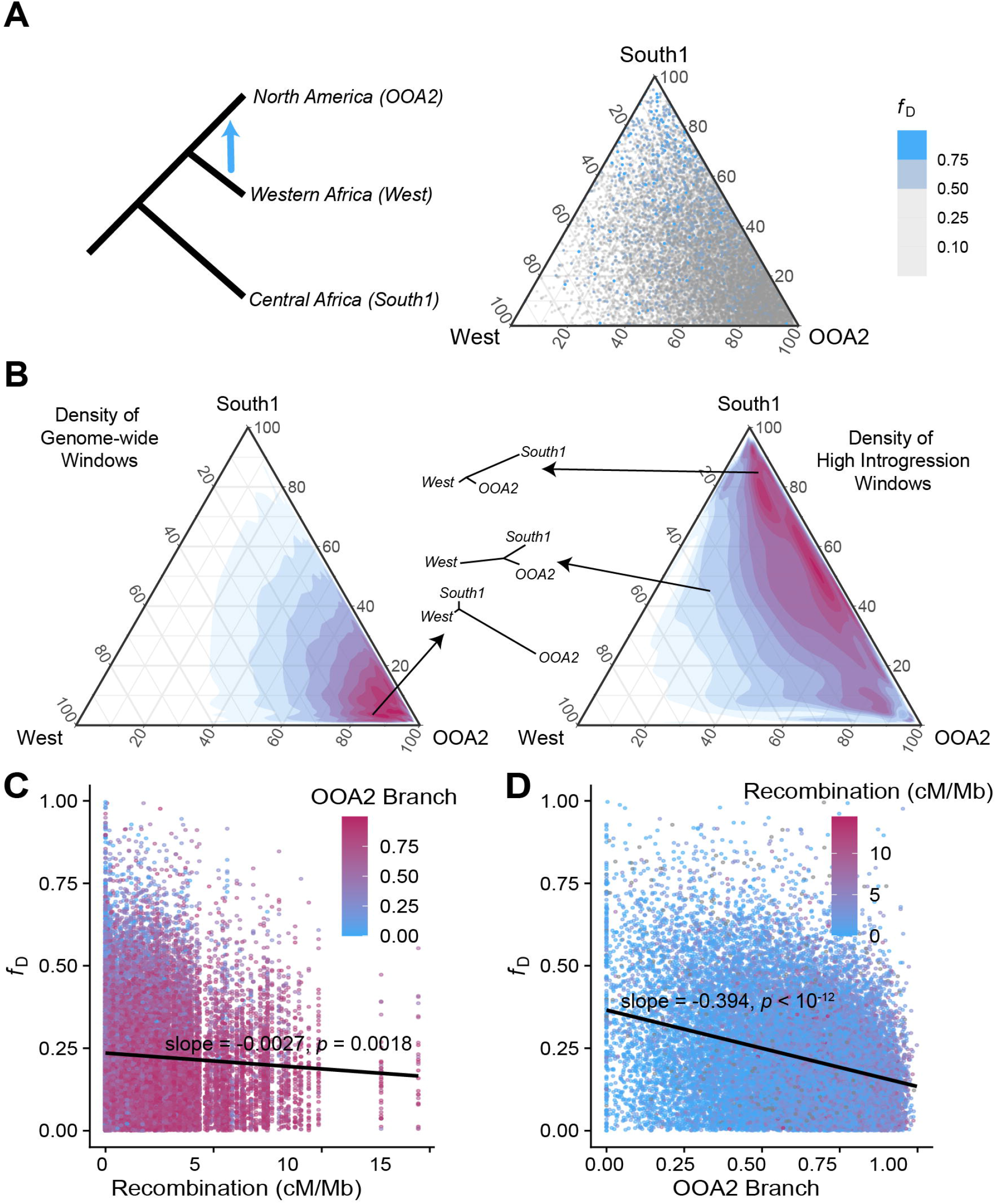
**(A)** Previous work has identified an introgression event between Western African *D. melanogaster* (West) into North American populations (OOA2). South African populations (South1) are ancestral to both and used as an outgroup. Using previously published data, we calculated relative branch scores (normalized PBS) for 25,874 windows across the genome with some evidence for introgression. Grey points indicate windows with *f*_D_ scores below 0.5, while blue points are windows with high evidence for introgression. **(B)** The genome-wide density of relative branch scores is clustered at values indicating high differentiation between OOA2 and both of the other populations (red values – high density, blue – low). However, the higher introgression windows (*f*_D_ > 0.5) are enriched for windows with longer West branches, as well as long South1 branches. The latter may indicate ILS or introgression. **(C)** A negative relationship between local recombination rate and introgression exists for these windows and **(D)** a similar negative relationship exists for OOA2 branch score and introgression, indicating diverged windows are more resilient to introgression. Slopes and *p*-values from a joint linear model with both terms modeled are listed on plots.

## Discussion

Our model describes the fitness of an introgressing haplotype over the course of divergence. Divergence, the fraction of the haplotype which carries replacement alleles, and (in case of diploidy) epistatic dominance play the largest roles in determining whether and what kinds of haplotypes can introgress, and we discuss each of these in turn.

### Divergence

Early in divergence, before species have evolved reproductive isolation, an introgressing haplotype is primarily contributing novel alleles. These alleles are likely to be neutral or positively selected to have been fixed in the source population. While these alleles may have negative epistatic interactions with alleles fixed in the receiving population, there are relatively few of these possible incompatibilities (Orr 1995; Turelli and Orr 2000). Additionally, there is likely positive epistasis between alleles the introgressing haplotype is carrying (Dagilis et al. 2019). The number of deleterious interactions with alleles in the receiving population increases linearly with the size of the introgressing haplotype, while the number of positive epistatic interactions increases at least quadratically (Equation 6). In turn, larger introgressing haplotypes are expected to, on average, be more positively selected. Even for alleles which are slightly deleterious in the host population, positive epistasis may allow larger haplotypes to introgress (Figure 2A). The same is largely true for haplotypes introgressing in a diploid population, however the dominance of epistasis plays a large role in determining selection on the introgressing haplotype (Figure S1). Gene flow between populations with low levels of divergence is not often dissected in the same way as gene flow between diverged species, but several studies do suggest that, in general, the fraction of the genome which shows evidence of introgression decreases with increasing divergence (Hamlin et al. 2020). Whether larger haplotypes are also introgressing at lower degrees of divergence is, as far as we are aware, unknown, but this pattern should in part result in a negative relationship between recombination rate and introgression, which has been observed in introgression between populations of *D. melanogaster* (Pool 2015; Duranton and Pool 2022), and is repeated in this study (Figure 4C).

As species continue to diverge (and *b* increases), there is a buildup of reproductive incompatibilities leading to increased selection against introgressing haplotypes. In turn, it takes larger introgressing haplotypes to overcome the deleterious effects with the host population, leading to an increasingly negative relationship between the size of the haplotype and its fitness (Figure 2A). Therefore, introgression of smaller haplotypes, which are on average more weakly selected against, is much more likely than introgression of large haplotypes. This relationship should manifest in a pattern of decreased introgression in regions of the genome with low recombination, a pattern observed in several species (Schumer et al. 2017). These results are also congruent with previous models which assume that larger introgressing haplotypes are broken down by recombination to avoid direct negative fitness effects (Harris and Nielsen 2016) or to break up negative epistatic interactions (Martin et al. 2019).

### Fraction of ancestral and replacement alleles

Not only is the haplotype bringing in novel alleles, which now have many potential incompatibility partners, but an introgressing haplotype is also likely to replace existing derived alleles in the host population (for instance, see Rinker et al. (2020)). Whether this replacement occurs in the form of reintroducing ancestral variants or “replacement” alleles (Figure 1B), larger haplotypes replace more diverged alleles, disrupting existing epistasis in the population. As we note earlier, this positive epistasis within the population has evolved under a selective sieve, and so it is likely to both be on average positive and of a larger magnitude than random new interactions between alleles untested in the same genetic background (Dagilis et al. 2019). Because of this, as larger haplotypes replace more local alleles they remove increasing numbers of positive interactions and may be strongly selected against (Figure 1C). This effect is paralleled in diploid populations (Supplementary Figure 3), although it depends strongly on the dominance of epistatic interactions. Whether the introduced alleles are simply bringing in ancestral variation or replacing existing derived variants with new ones, haplotypes carrying many such alleles are always more strongly selected against than ones carrying purely novel variants (Figure 3). This means that introgression is only expected to be resisted in parts of the genome which have large numbers of diverged alleles, but not in regions carrying primarily ancestral variants. We find this pattern among populations of *D. melanogaster*, with both a negative relationship between introgression and divergence of the host population (OOA2) from the source (West) (Figure 4D) as well as an over-representation of windows in which the West population shows rapid evolution among the introgressed regions (Figure 4B).

### Dominance of epistasis

In the haploid case, we can infer some reasonable parameter values (specifically for epistasis within and between populations) to make some general conclusions. Such conclusions are difficult to extend to diploids without a clear understanding of the dominance of epistatic interactions. While rare, the introgressing haplotype will occur primarily in heterozygotes. As such, when epistasis is recessive, introgressing haplotypes will not be benefitting from positive interactions among the alleles they carry, but replacement alleles will strongly disrupt existing positive epistasis, leading to overall stronger selection against introgression (Figure S1). On the other hand, if epistasis is dominant, introgressing haplotypes do not disrupt existing interactions, and will bring in stronger positive effects among the alleles they carry, leading to much stronger selection for introgressed haplotypes of any size. Little is known about the dominance of epistatic interactions, but they may be recessive as suggested by studies of the large-X effect (Turelli and Orr 2000) and Haldane’s Rule (Coyne and Orr 1997). As with any biological system, in truth substitutions are likely to have epistatic interactions of various dominance, and so a single parameter for dominance is unlikely to fully represent reality. Nonetheless, the general shape of the fitness functions does not change based on the dominance parameters examined here, and so most of our predictions should hold in a variety of circumstances.

### Caveats

Our model is simplistic by design, in order to give a clearer intuition of what the fitness of an introgressing haplotype of any size during the course of divergence is. To be able to ask this question, we make several major assumptions. First, we assume that the haplotype has appeared in the host population, but do not know how. In order for a small haplotype to occur in the host population, many generations of backcrossing must first occur. Later in divergence, when selection against large haplotypes is very strong, it may be exceedingly rare for these small haplotypes to make it across species barriers, primarily because early generation backcrosses are strongly selected against (see Matute and Cooper (2021) for recent reviews). Indeed, we do not account for the fitness of F1 or later generation hybrids at all and simply measure the fitness of the haplotype in an otherwise uniform genetic background.

Second, we ignore recombination in our model, which prompts several important caveats. The haplotype’s fitness determines its fixation only if that haplotype stays the same size. The positive fitness effects experienced by large haplotypes early in divergence could in practice be much weaker, as recombination breaks these haplotypes up. It is possible that the haplotype is being maintained at this size by an inversion or some recombination modifier linked to the haplotype, acting as a “supergene” (Thompson and Jiggins 2014; Kirubakaran et al. 2016; Jay et al. 2018; Kim et al. 2022), but most introgressing regions are unlikely to be maintained in perfect LD. Similarly, because we ignore recombination we equate the “size” of the haplotype to the numbers of alleles it carries with its physical size. However, in practice, physically small regions may carry many more alleles than large ones. As a result, our expectation of different relationships between recombination rate and introgression probability depends in large part on an assumption of a roughly equal density of substitutions across the genome.

Lastly, we assume that there are only two genotypes in the population – individuals carrying the “host” genetic background and those carrying the introgressing haplotype. Even if we ignore polymorphism within the host population before hybridization, which may be highly relevant to fitness of hybrids in general (Cutter 2012; Maya-Lastra and Eaton 2021), the two processes above – backcrossing and recombination, will lead to a wide array of genotypes in the host population. Real instances of introgression have the haplotypes competing not solely against a host genotype, but also against many other introgressing haplotypes of various sizes and distributions along the genome. Early in divergence, this may dampen the benefits of keeping large blocks intact. More broadly, this means that multiple haplotypes of varying sizes are likely to be introgressing, making predictions far less straightforward (Veller et al. 2021; Duranton and Pool 2022; Pfennig and Lachance 2022).

### Conclusions

Selection for or against introgressed haplotypes seems to follow similar patterns to fitness of hybrids in general. Early in divergence, hybrid fitness can remain high (Simon et al. 2018; Dagilis et al. 2019), and we find that fitness changes exponentially with respect to the size of the introgressing haplotype. Under reasonable parameters, this leads to positive selection of large introgressing haplotypes early in divergence, especially when they carry alleles at sites that have not experienced substitutions in the host population. On the other hand, later in divergence, as hybrid fitness decreases, haplotypes are negatively selected unless they are bringing in vast numbers of novel alleles, which is unlikely to occur in nature. Therefore, different relationships between introgressing haplotype size and fitness are expected between recently diverged populations and highly diverged species pairs.

There are two consequences from these simple observations. Late in divergence, recombination that breaks up introgressing haplotypes is nearly always favored, as smaller haplotypes are less strongly selected against. This expectation may explain the observed relationship between introgression and recombination in several systems, including swordtail fish, *Heliconius* butterflies and Neanderthal-human introgression (Schumer et al. 2017; Martin et al. 2019). However, early in divergence, larger haplotypes might actually introgress more easily. In turn, the relationship between recombination rate and introgression may not exist, or even be reversed. The second major result is that introgression should be impacted by how many diverged alleles an introgressing haplotype is replacing. Introgression of a haplotype that carries almost entirely new derived alleles is almost always more beneficial than a haplotype replacing existing derived alleles in the population (see Figure 2B, Figure 3).

The predictions of our model generate several major expectations for patterns of introgression in nature that we cannot test here. First, large haplotypes that have managed to introgress should carry proportionately few ancestral or replacement alleles, as they should be strongly selected against otherwise. Second, there should be an asymmetry in the direction of introgression, with species that are more highly diverged in the pair showing less evidence of introgression. Haplotypes introgressing into a more diverged population are more likely to “overwrite” existing derived alleles, and so are more strongly selected against than haplotypes introgressing into a genetic background that has fewer substitutions in general. A rapidly adapting population therefore ends up with a genome that is more resilient to introgression than a sister population drifting around the ancestral optimum. Finally, regions of the genome which have many fixed substitutions may be resilient to introgression. However, divergence at a single population is insufficient to generate an “island of speciation” (Turner et al. 2005; Turner and Hahn 2010; Cruickshank and Hahn 2014), as introgression into the population carrying largely ancestral variants is still possible. Instead, islands of speciation that are resilient to introgression between populations will only be generated when both populations fix multiple substitutions within the same region.

The current model shows how basic intuitions about the size of introgressing blocks are affected by epistatic fitness effects on the block of introgressing alleles and with the host genome. The results we obtain depend on several parameters that require far more study. Very few studies to date have examined the strength of epistatic interactions, and so parameterizing our model is still difficult. Similarly, very little is known about the dominance of epistasis. Lastly, our model, like any model, makes simplifying assumptions that are not likely to hold in many natural cases of introgression. Nonetheless, this work provides a novel framework to understand the fitness effects caused by the interplay between the size of introgressing allele blocks and divergence time.

## Acknowledgements

We would like to thank Adam Stuckert, Jenn Coughlan, Gaston Jofre, Jonathan Rader and Sean Anderson for helpful comments. This work was supported by the National Institute of General Medical Sciences of the National Institutes of Health (NIH) under Award R01GM125715 to DRM. AJD was supported under the National Institute of Allergy and Infectious Diseases of the National Institutes of Health (NIH) Award T32-AI052080.

## Materials and methods and model details

We model the fitness of an introgressing haplotype that carries *x* alleles that are not present in the host population. For simplicity, we assume that the host population does not carry any polymorphisms aside from the introgressing haplotype. We will only examine the fitness of the introgressing haplotype while it is rare. As such, we assume that the average fitness of the population is determined entirely by the fitness of locally fixed alleles, relative to the fitness of a purely ancestral genome set to 1. The fitness of an individual carrying a set of derived alleles X is defined to be:

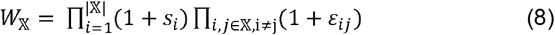

Where *s*_*x*_ is the direct selection coefficient of allele *x* and *ε*_*ij*_ is the epistatic effect between alleles *i* and *j*. That is, we assume that fitness effects are multiplicative, although largely the same results are obtained assuming additive fitness effects (results not shown). The fitness of an individual carrying the introgressing haplotype will therefore depend on the alleles the haplotype is carrying – these will be the diverged alleles outside the haplotype (𝔹\𝕏 _B_, i.e. all diverged alleles in B minus those that are overwritten by the introgressing haplotype) and the new diverged alleles brought in by the haplotype (set (𝕏_A_). A carrier of an introgressed haplotype therefore has the set of alleles □=𝕏_*A*_U (𝔹\𝕏_B_. The fitness of an individual carrying the introgressed haplotype is therefore:

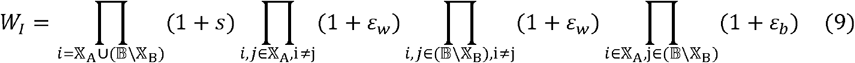

Where *ε*_*W*_ is the average epistatic effect for alleles fixed “within” the same population, while ε_*b*_ is the epistatic effect for alleles fixed “between” populations. The relative selection coefficient can then be calculated as *W*_I_ divided by the average fitness of an individual in B, given by the left side of Equation (1), minus 1. These equations are used to produce Figure 2. To obtain analytically simple results, we note that when the epistatic effects and direct selection are very small, the above is well approximated by:

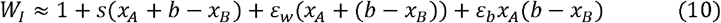

Where *b* = |𝔹|, or the size of set 𝔹. Note that alleles in sets 𝕏_A_ and 𝕏_A_ can overlap (i.e. a replacement allele is a locus where population B has a fixed substitution that also has a fixed substitution in population A). If we think of the three types of introgressing alleles outlined in Figure 1B, *x*_A_ = *x*_new_ + *x*_replacement_, while *x*_B_ = *x*_ancestral_ + *x*_replacemen**t**._ Early in the process of divergence, it’s reasonable to assume that the number of replacement alleles will be negligible, and so to simplify results we assume that *x*_replacement_ = 0, but see Figure 3 for an exploration of how replacement alleles affect the fitness of the introgressing haplotype. We can next assume that *x*_A_ = (1-*f*)*x* and *x*_B_ = *f x*, and plug in these values to obtain Equation (6). Finally, we can take the derivative of the resulting equation with respect to *x* while setting *f*=½ to obtain Equation (7).

### Diploid model

We now need to consider the diploid case. We again assume the haplotype is quite rare, thus will always be found in a heterozygous state. As such, we need to consider two types of dominance effects - dominance of direct and epistatic effects. We assume direct effects have, on average, dominance *h*. Epistasis between two heterozygous derived alleles will be assumed to be of strength *a*_1_*ε*, while epistasis between a homozygous and heterozygous derived pair of alleles is of strength *a*_2_*ε*. Epistasis between two homozygous derived alleles is of full strength E. Like the haploid model, we can express the fitness of an individual carrying the introgressed haplotype as follows:

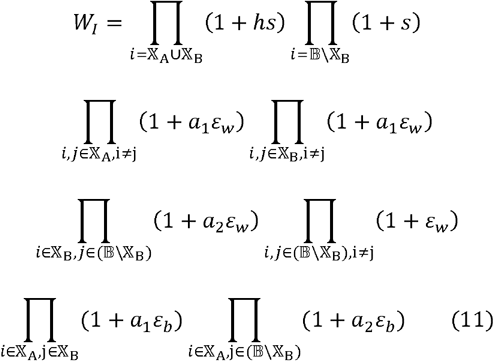

And the effective selection coefficient can be calculated by comparing this fitness to the fitness of an average individual in the population, still equal to Equation (1). These equations are used to produce Supplementary Figure 3. Following the same logic as before, we can approximate the relative selection coefficient further by assuming all effects are small and have low variance to produce:

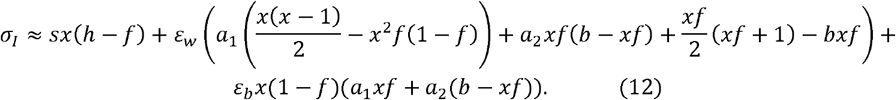

Finally, we can simplify the above in the special case that epistasis is additive (*a*_1_= ½ *a*_2_ = ¼), to obtain a slightly condensed form:

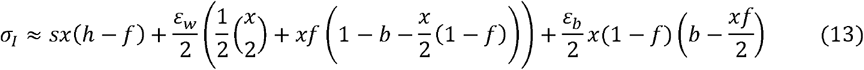

A few takeaways can be obtained from these equations. Just as in the haploid case, a linear relationship between the effective selection coefficient and divergence is expected. On the other hand, the size of the haplotype appears primarily in quadratic terms, and so large haplotypes at low divergence may once again be positively selected fairly easily. Finally, we can make a note on dominance – if direct selection is on average additive (*h* = ½) and the introgressing haplotype carries new alleles as frequently as ancestral (*f*= ½), no net direct fitness effects are expected and only epistatic effects determine selection on introgressing haplotypes.

### Re-analysis of *Drosophila melanogaster* data

We used previously calculated values of *f*_D_ for introgression between fly lines with a majority ancestry West and those with majority ancestry OOA2 (Coughlan et al. 2021). Individuals with less than 1% introgressed ancestry, but majority of either OOA2, West, or South1 ancestry were identified, and *F*_ST_ between these individuals was calculated using *pixy* (Korunes and Samuk 2021) in windows matching the windows from introgression analyses. PBS (Yi et al. 2010) values for each branch were next calculated. Windows with negative PBS values for any of the windows were next filtered out, however major results are largely the same if these windows are kept (Supplementary Figure 4). The resulting values were plotted in ternary plots using *ggplot* (Wickham 2016) and *ggtern* (Hamilton and Ferry 2018) packages. Recombination rates per window were obtained using data from (Comeron et al. 2012), assigning average recombination rate for each window based on the Comeron dataset. We next fit a general linear model of *f*_D_ dependent on recombination and PBS_OOA2_. While we could include other PBS branches as well, their high degree of correlation made us cautious of using more than one PBS branch in the model at a time.

## Supplementary Figures

**Supplementary Figure 1.**
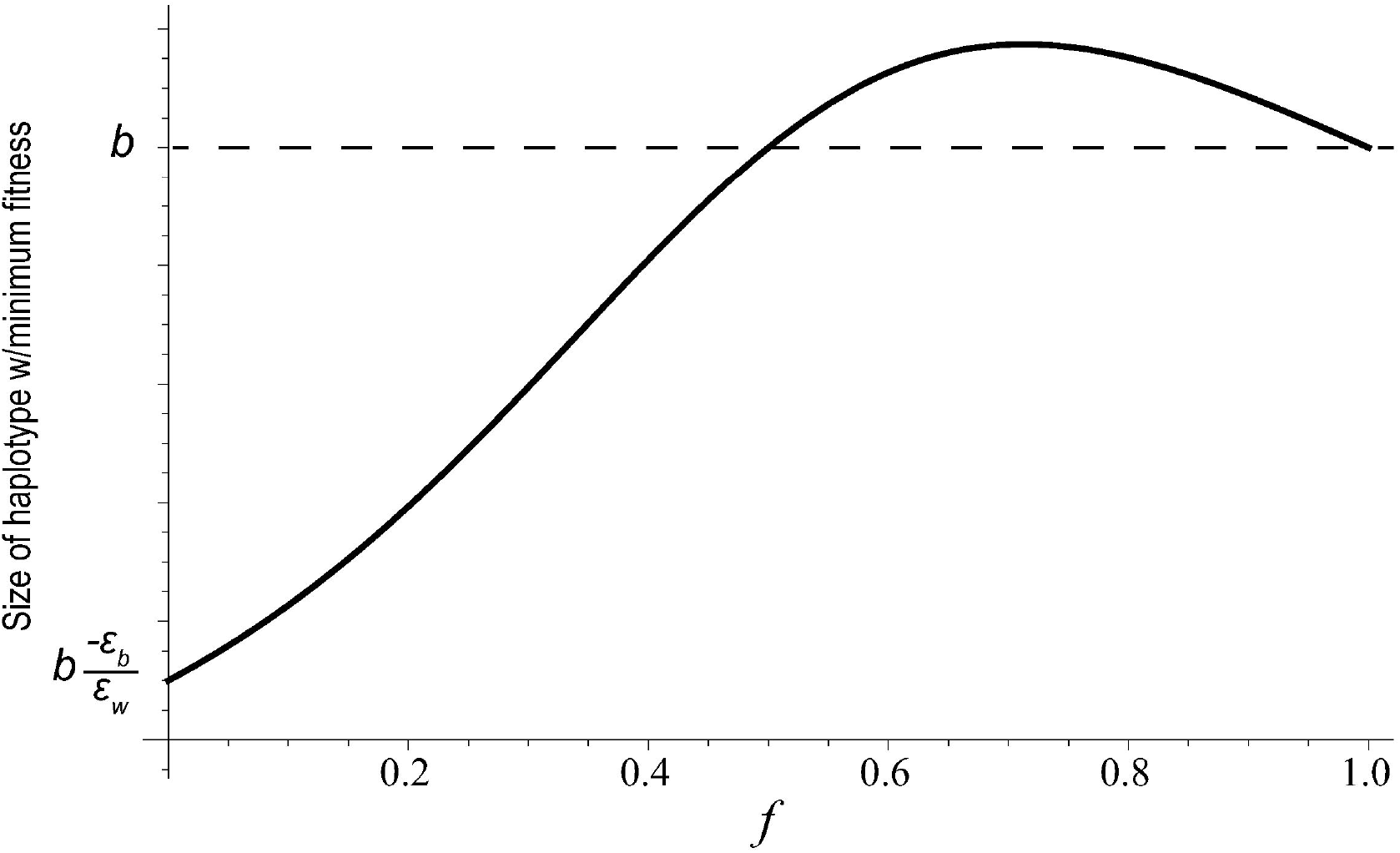
A graphical illustration of Equation 7, demonstrating the size at which the introgressing haplotype is most strongly selected against versus the fraction of the haplotype that carries ancestral/replacement alleles. The above shape is consistent for all values of parameters, and values are all positive when between and within population epistasis are opposite signs or are both positive.

**Supplementary Figure 2:**
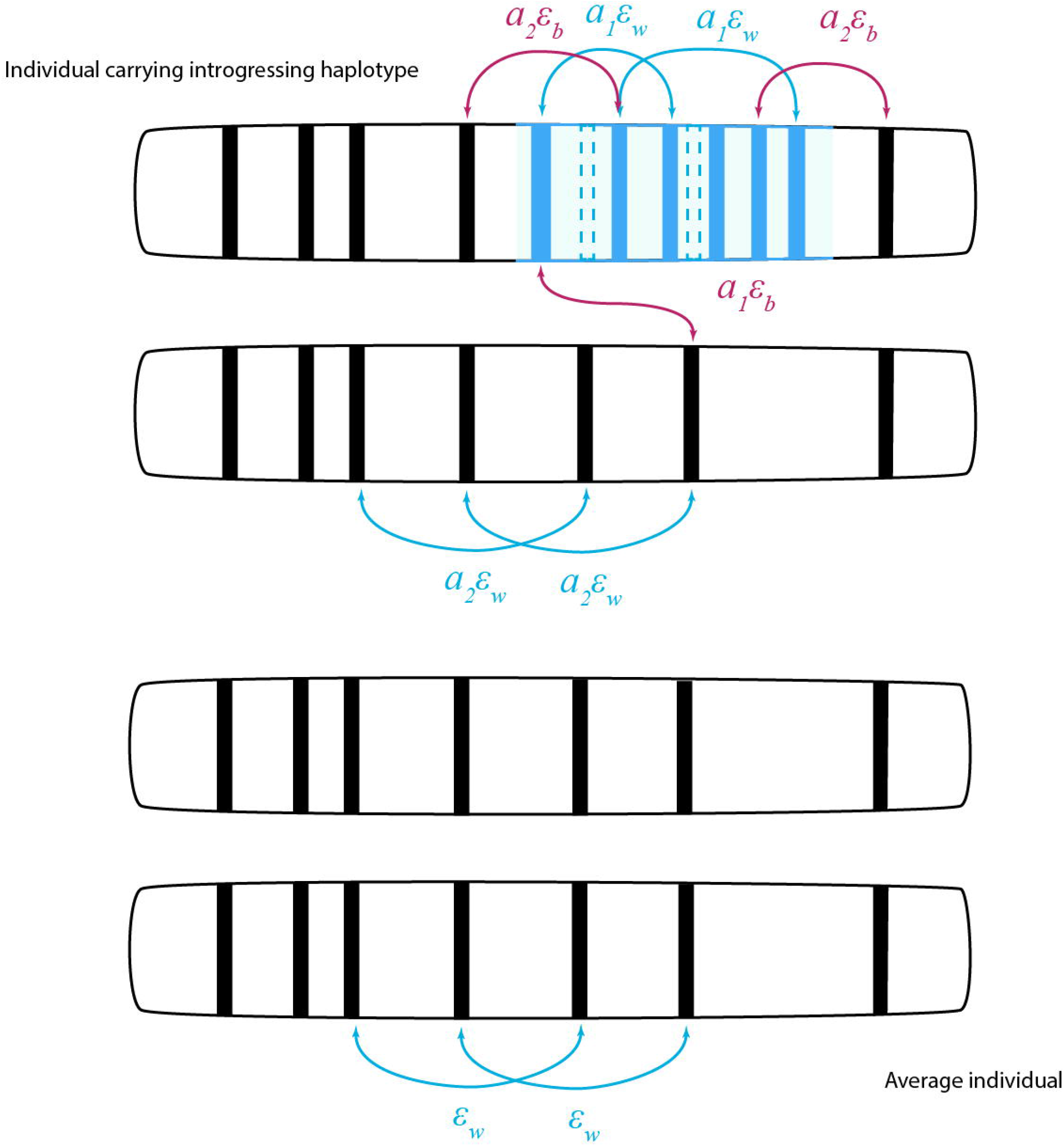
Schematic representation of fitness effects in a diploid model. Individuals carrying the introgressed haplotype are assumed to always be heterozygous, therefore carrying epistatic effects of varying degrees of dominance, with all new within population epistatic interactions occurring between heterozygous alleles, while some between population interactions occur between heterozygous and homozygous alleles.

**Supplementary Figure 3.**
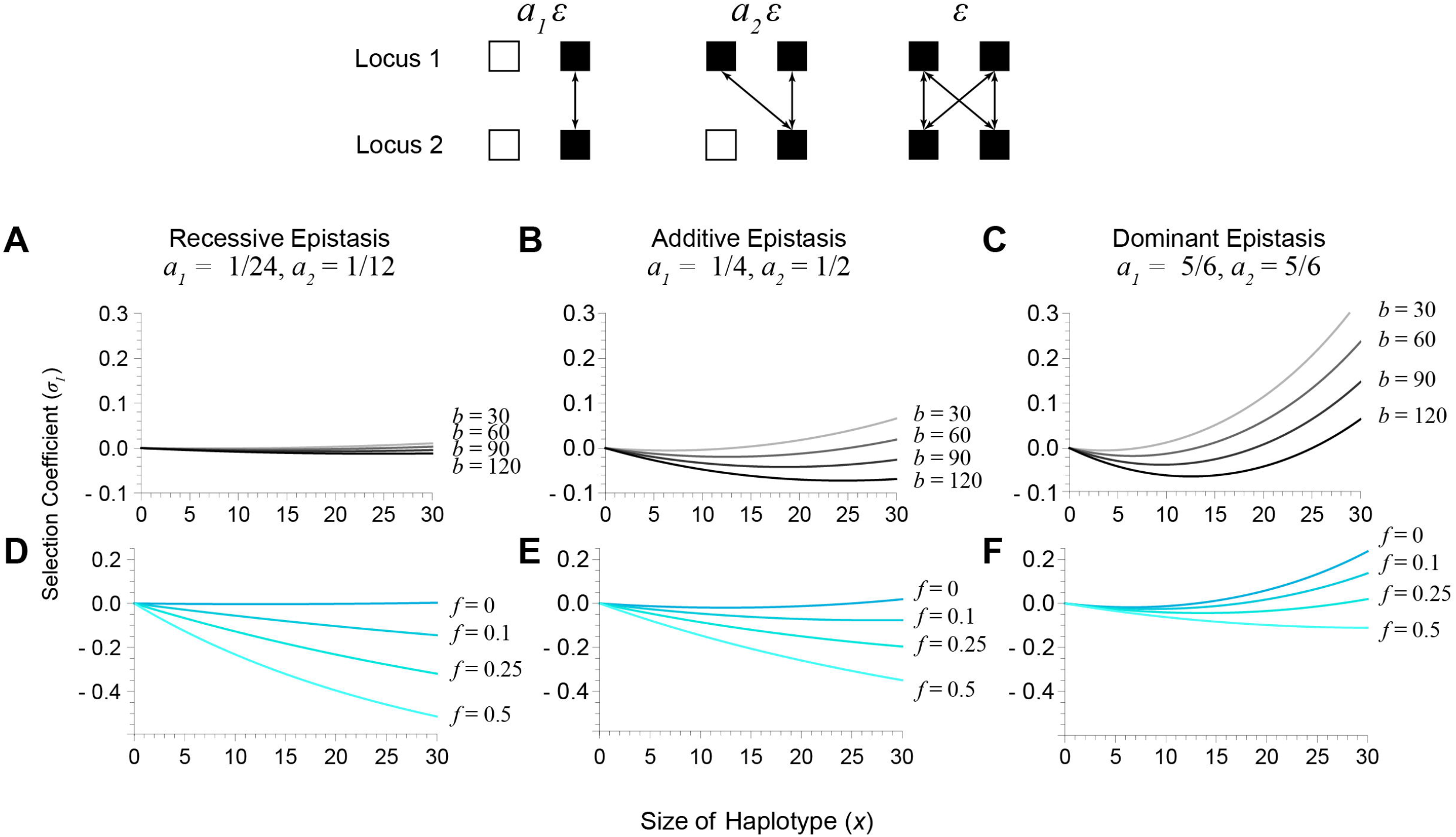
Results for a diploid model. All parameters are the same to those used in Figure 2, and only exact numeric solutions are shown. Two new parameters, *a*_1_ and *a*_2_ determine the dominance of epistasis. When an individual is heterozygous at two derived alleles, epistasis is of strength *a*_1_* *ε*, when one is heterozygous and the other is homozygous it is *a*_2_\**ε*, and when both are homozygous it is epsilon. Top row of plots shows how varying divergence (b) changes selection on introgressing haplotypes of varying sizes, *f* fixed at 0. Bottom row shows the effects of varying the fraction of the haplotype that is replacement alleles (f), *b* fixed at 60. Patterns in general mirror the haploid case, although recessive epistasis leads to much stronger selection against the introgressing haplotype, and dominant epistasis leads to overall stronger selection for introgression. As with other models where *f* is used, we assume that the number of replacement alleles is 0, but the shape of the curve changes very little with the addition of replacement alleles (as each replacement allele increases *x* by only 1, but carries the effects of both a new and ancestral alleles).

**Supplementary Figure 4.**
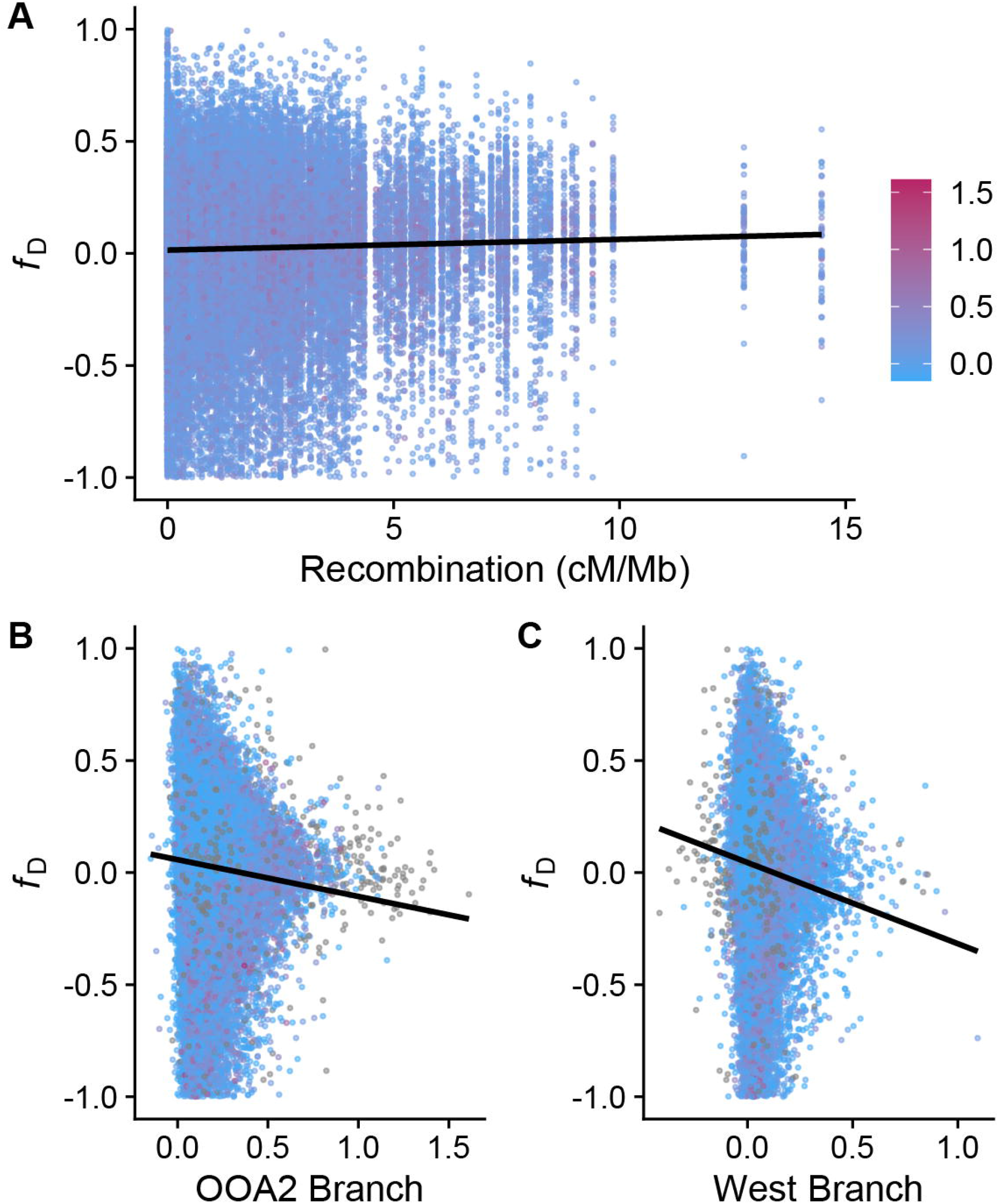
Inclusion of negative *f*_D_ values (support for introgression in the opposite biologically plausible direction) and negative PBS. (A) A positive relationship between recombination and *f*_D_ is found, but is driven primarily by large numbers of negative *f*_D_ values at low recombination windows. Nonetheless, a negative relationship between branch length and *f*_D_ value is found for both OOA2 PBS values and West PBS values (B and C).

## References

Barton, N. H. 2001. The role of hybridization in evolution. Mol. Ecol. 10:551–568.

Barton, N. H. and B. O. Bengtsson. 1986. The barrier to genetic exchange between hybridising populations. Heredity (Edinb.) 57:357–376.

Comeron, J. M., R. Ratnappan, and S. Bailin. 2012. The many landscapes of recombination in Drosophila melanogaster. PLoS Genet. 8:e1002905.

Costanzo, M., B. VanderSluis, E. N. Koch, A. Baryshnikova, C. Pons, G. Tan, W. Wang, M. Usaj, J. Hanchard, S. D. Lee, V. Pelechano, E. B. Styles, M. Billmann, J. van Leeuwen, N. van Dyk, Z. Y. Lin, E. Kuzmin, J. Nelson, J. S. Piotrowski, T. Srikumar, S. Bahr, Y. Chen, R. Deshpande, C. F. Kurat, S. C. Li, Z. Li, M. M. Usaj, H. Okada, N. Pascoe, B. J. San Luis, S. Sharifpoor, E. Shuteriqi, S. W. Simpkins, J. Snider, H. G. Suresh, Y. Tan, H. Zhu, N. Malod-Dognin, V. Janjic, N. Przulj, O. G. Troyanskaya, I. Stagljar, T. Xia, Y. Ohya, A. C. Gingras, B. Raught, M. Boutros, L. M. Steinmetz, C. L. Moore, A. P. Rosebrock, A. A. Caudy, C. L. Myers, B. Andrews, and C. Boone. 2016. A global genetic interaction network maps a wiring diagram of cellular function. Science 353.

Coughlan, J. M., A. J. Dagilis, A. Serrato-Capuchina, H. Elias, D. Peede, K. Isbell, D. M. Castillo, B. S. Cooper, and D. R. Matute. 2021. Patterns of and processes shaping population structure and introgression among recently differentiated Drosophila melanogaster populations. bioRxiv:2021.2006.2025.449842.

Coughlan, J. M. and D. R. Matute. 2020. The importance of intrinsic postzygotic barriers throughout the speciation process. Philosophical Transactions of the Royal Society B: Biological Sciences 375:20190533.

Coyne, J. A. and H. A. Orr. 1997. “Patterns of speciation in Drosophila” revisited. Evolution 51:295–303.

Coyne, J. A. and H. A. Orr. 2004. Speciation. Sinauer Associates, Sunderland, MA.

Cruickshank, T. E. and M. W. Hahn. 2014. Reanalysis suggests that genomic islands of speciation are due to reduced diversity, not reduced gene flow. Mol. Ecol. 23:3133–3157.

Cutter, A. D. 2012. The polymorphic prelude to Bateson–Dobzhansky–Muller incompatibilities. Trends Ecol. Evol. 27:209–218.

Dagilis, A. J., M. Kirkpatrick, and D. I. Bolnick. 2019. The evolution of hybrid fitness during speciation. PLoS Genet. 15:e1008125.

Dagilis, A. J., D. Peede, J. M. Coughlan, G. I. Jofre, E. R. R. D’Agostino, H. Mavengere, A. D. Tate, and D. R. Matute. 2022. A need for standardized reporting of introgression: Insights from studies across eukaryotes. Evol. Letters n/a.

Dixon, G., J. Kitano, and M. Kirkpatrick. 2019. The origin of a new sex chromosome by introgression between two stickleback fishes. Mol. Biol. Evol. 36:28–38.

Duranton, M. and J. E. Pool. 2022. Interactions Between Natural Selection and Recombination Shape the Genomic Landscape of Introgression. Mol. Biol. Evol. 39.

Edelman, N. B., P. B. Frandsen, M. Miyagi, B. Clavijo, J. Davey, R. B. Dikow, G. Garcia-Accinelli, S. M. Van Belleghem, N. Patterson, D. E. Neafsey, R. Challis, S. Kumar, G. R. P. Moreira, C. Salazar, M. Chouteau, B. A. Counterman, R. Papa, M. Blaxter, R. D. Reed, K. K. Dasmahapatra, M. Kronforst, M. Joron, C. D. Jiggins, W. O. McMillan, F. Di Palma, A. J. Blumberg, J. Wakeley, D. Jaffe, and J. Mallet. 2019. Genomic architecture and introgression shape a butterfly radiation. Science 366:594–599.

Edelman, N. B. and J. Mallet. 2021. Prevalence and adaptive impact of introgression. Annu. Rev. Genet. 55:265–283.

Fontaine, M. C., J. B. Pease, A. Steele, R. M. Waterhouse, D. E. Neafsey, I. V. Sharakhov, X. Jiang, A. B. Hall, F. Catteruccia, E. Kakani, S. N. Mitchell, Y. C. Wu, H. A. Smith, R. R. Love, M. K. Lawniczak, M. A. Slotman, S. J. Emrich, M. W. Hahn, and N. J. Besansky. 2015. Extensive introgression in a malaria vector species complex revealed by phylogenomics. Science 347:1258524.

Fraïsse, C., P. A. Gunnarsson, D. Roze, N. Bierne, and J. J. Welch. 2016. The genetics of speciation: insights from Fisher’s geometric model. Evolution 70:1450–1464.

Gavrilets, S. 2004. Fitness landscapes and the origin of species (MPB-41). Princeton University Press.

Hamilton, N. E. and M. Ferry. 2018. ggtern: Ternary Diagrams Using ggplot2. Journal of Statistical Software, Code Snippets 87:1 – 17.

Hamlin, J. A. P., M. S. Hibbins, and L. C. Moyle. 2020. Assessing biological factors affecting postspeciation introgression. Evol. Letters 4:137–154.

Harris, K. and R. Nielsen. 2016. The Genetic Cost of Neanderthal Introgression. Genetics 203:881–891.

Huerta-Sánchez, E., X. Jin, Asan, Z. Bianba, B. M. Peter, N. Vinckenbosch, Y. Liang, X. Yi, M. He, M. Somel, P. Ni, B. Wang, X. Ou, Huasang, J. Luosang, Z. X. P. Cuo, K. Li, G. Gao, Y. Yin, W. Wang, X. Zhang, X. Xu, H. Yang, Y. Li, J. Wang, J. Wang, and R. Nielsen. 2014. Altitude adaptation in Tibetans caused by introgression of Denisovan-like DNA. Nature 512:194–197.

Jay, P., A. Whibley, L. Frézal, M. Á. Rodríguez de Cara, R. W. Nowell, J. Mallet, K. K. Dasmahapatra, and M. Joron. 2018. Supergene Evolution Triggered by the Introgression of a Chromosomal Inversion. Curr. Biol. 28:1839-1845.e1833.

Juric, I., S. Aeschbacher, and G. Coop. 2016. The Strength of Selection against Neanderthal Introgression. PLoS Genet. 12:e1006340.

Kim, K.-W., R. De-Kayne, I. J. Gordon, K. S. Omufwoko, D. J. Martins, R. ffrench-Constant, and S. H. Martin. 2022. Stepwise evolution of a butterfly supergene via duplication and inversion. Philosophical Transactions of the Royal Society B: Biological Sciences 377:20210207.

Kirubakaran, T. G., H. Grove, M. P. Kent, S. R. Sandve, M. Baranski, T. Nome, M. C. De Rosa, B. Righino, T. Johansen, H. Ottera, A. Sonesson, S. Lien, and O. Andersen. 2016. Two adjacent inversions maintain genomic differentiation between migratory and stationary ecotypes of Atlantic cod. Mol. Ecol. 25:2130–2143.

Korunes, K. L. and K. Samuk. 2021. pixy: Unbiased estimation of nucleotide diversity and divergence in the presence of missing data. Mol. Ecol. Resour. 21:1359–1368.

Martin, S. H., J. W. Davey, C. Salazar, and C. D. Jiggins. 2019. Recombination rate variation shapes barriers to introgression across butterfly genomes. PLoS Biol. 17:e2006288.

Martin, S. H. and C. D. Jiggins. 2017. Interpreting the genomic landscape of introgression. Curr. Opin. Genet. Dev. 47:69–74.

Matute, D. R., I. A. Butler, D. A. Turissini, and J. A. Coyne. 2010. A test of the snowball theory for the rate of evolution of hybrid incompatibilities. Science 329:1518–1521.

Matute, D. R. and B. S. Cooper. 2021. Comparative studies on speciation: 30 years since Coyne and Orr. Evolution 75:764–778.

Maya-Lastra, C. A. and D. A. R. Eaton. 2021. Genetic incompatibilities do not snowball in a demographic model of speciation. bioRxiv:2021.2002.2023.432472.

Meisner, J. and A. Albrechtsen. 2018. Inferring Population Structure and Admixture Proportions in Low-Depth NGS Data. Genetics 210:719–731.

Moyle, L. C. and T. Nakazato. 2010. Hybrid incompatibility “snowballs” between Solanum species. Science 329:1521–1523.

Muirhead, C. A. and D. C. Presgraves. 2016. Hybrid Incompatibilities, Local Adaptation, and the Genomic Distribution of Natural Introgression between Species. Am. Nat. 187:249–261.

Norris, L. C., B. J. Main, Y. Lee, T. C. Collier, A. Fofana, A. J. Cornel, and G. C. Lanzaro. 2015. Adaptive introgression in an African malaria mosquito coincident with the increased usage of insecticide-treated bed nets. Proceedings of the National Academy of Sciences 112:815–820.

Orr, H. A. 1995. The population genetics of speciation: the evolution of hybrid incompatibilities. Genetics 139:1805–1813.

Petr, M., S. Pääbo, J. Kelso, and B. Vernot. 2019. Limits of long-term selection against Neandertal introgression. Proceedings of the National Academy of Sciences 116:1639–1644.

Pfennig, A. and J. Lachance. 2022. Hybrid fitness effects modify fixation probabilities of introgressed alleles. G3 Genes|Genomes|Genetics 12.

Poelstra, J. W., N. Vijay, C. M. Bossu, H. Lantz, B. Ryll, I. Muller, V. Baglione, P. Unneberg, M. Wikelski, M. G. Grabherr, and J. B. Wolf. 2014. The genomic landscape underlying phenotypic integrity in the face of gene flow in crows. Science 344:1410–1414.

Pool, J. E. 2015. The Mosaic Ancestry of the Drosophila Genetic Reference Panel and the D. melanogaster Reference Genome Reveals a Network of Epistatic Fitness Interactions. Mol. Biol. Evol. 32:3236–3251.

Ravinet, M., K. Yoshida, S. Shigenobu, A. Toyoda, A. Fujiyama, and J. Kitano. 2018. The genomic landscape at a late stage of stickleback speciation: High genomic divergence interspersed by small localized regions of introgression. PLoS Genet. 14:e1007358.

Rinker, D. C., C. N. Simonti, E. McArthur, D. Shaw, E. Hodges, and J. A. Capra. 2020. Neanderthal introgression reintroduced functional ancestral alleles lost in Eurasian populations. Nat. Ecol. Evol. 4:1332–1341.

Roux, C., C. Fraisse, J. Romiguier, Y. Anciaux, N. Galtier, and N. Bierne. 2016. Shedding Light on the Grey Zone of Speciation along a Continuum of Genomic Divergence. PLoS Biol. 14:e2000234.

Schumer, M., R. Cui, D. L. Powell, G. G. Rosenthal, and P. Andolfatto. 2016. Ancient hybridization and genomic stabilization in a swordtail fish. Mol. Ecol. 25:2661–2679.

Schumer, M., C. Xu, D. L. Powell, A. Durvasula, L. Skov, C. Holland, S. Sankararaman, P. Andolfatto, G. G. Rosenthal, and M. Przeworski. 2017. Natural selection interacts with the local recombination rate to shape the evolution of hybrid genomes. bioRxiv:212407.

Silva, D. N., V. Varzea, O. S. Paulo, and D. Batista. 2018. Population genomic footprints of host adaptation, introgression and recombination in coffee leaf rust. Mol Plant Pathol 19:1742–1753.

Simon, A., N. Bierne, and J. J. Welch. 2018. Coadapted genomes and selection on hybrids: Fisher’s geometric model explains a variety of empirical patterns. Evol Lett 2:472–498.

Thompson, M. J. and C. D. Jiggins. 2014. Supergenes and their role in evolution. Heredity (Edinb.) 113:1–8.

Turelli, M. and H. A. Orr. 2000. Dominance, epistasis and the genetics of postzygotic isolation. Genetics 154:1663–1679.

Turner, T. L. and M. W. Hahn. 2010. Genomic islands of speciation or genomic islands and speciation? Pp. 848--850.

Turner, T. L., M. W. Hahn, and S. V. Nuzhdin. 2005. Genomic islands of speciation in Anopheles gambiae. PLoS Biol. 3:e285.

Veller, C., N. B. Edelman, P. Muralidhar, and M. A. Nowak. 2021. Recombination and selection against introgressed DNA. bioRxiv:846147.

Wickham, H. 2016. ggplot2: elegant graphics for data analysis. Springer.

Yi, X., Y. Liang, E. Huerta-Sanchez, X. Jin, Z. X. P. Cuo, J. E. Pool, X. Xu, H. Jiang, N. Vinckenbosch, T. S. Korneliussen, H. Zheng, T. Liu, W. He, K. Li, R. Luo, X. Nie, H. Wu, M. Zhao, H. Cao, J. Zou, Y. Shan, S. Li, Q. Yang, Asan, P. Ni, G. Tian, J. Xu, X. Liu, T. Jiang, R. Wu, G. Zhou, M. Tang, J. Qin, T. Wang, S. Feng, G. Li, Huasang, J. Luosang, W. Wang, F. Chen, Y. Wang, X. Zheng, Z. Li, Z. Bianba, G. Yang, X. Wang, S. Tang, G. Gao, Y. Chen, Z. Luo, L. Gusang, Z. Cao, Q. Zhang, W. Ouyang, X. Ren, H. Liang, H. Zheng, Y. Huang, J. Li, L. Bolund, K. Kristiansen, Y. Li, Y. Zhang, X. Zhang, R. Li, S. Li, H. Yang, R. Nielsen, J. Wang, and J. Wang. 2010. Sequencing of 50 Human Exomes Reveals Adaptation to High Altitude. Science 329:75–78.

